# Probing and predicting ganglion cell responses to smooth electrical stimulation in healthy and blind mouse retina

**DOI:** 10.1101/609826

**Authors:** Larissa Höfling, Philipp Berens, Günther Zeck

## Abstract

Retinal implants are used to replace lost photoreceptors in blind patients suffering from retinopathies such as retinitis pigmentosa. Patients wearing implants regain some rudimentary visual function. However, it is severely limited compared to normal vision because non-physiological stimulation strategies fail to selectively activate different retinal pathways at sufficient spatial and temporal resolution. The development of improved stimulation strategies is rendered difficult by the large space of potential stimuli. Here we systematically explore a subspace of potential stimuli by electrically stimulating healthy and blind mouse retina in epiretinal configuration using smooth Gaussian white noise delivered by a high-density CMOS-based microelectrode array. We identify linear filters of retinal ganglion cells (RGCs) by fitting a linear-nonlinear-Poisson (LNP) model. Our stimulus evokes fast, reliable, and spatially confined spiking responses in RGC which are accurately predicted by the LNP model. Furthermore, we find diverse shapes of linear filters in the linear stage of the model, suggesting diverse preferred electrical stimuli of RGCs. Our smooth electrical stimulus could provide a starting point of a model-guided search for improved stimuli for retinal prosthetics.

## Introduction

Sensory neural prostheses attempt to replace nonfunctional elements of sensory organs and thereby to restore access to sensory information. For example, cochlear implants, which have been in clinical use for many years, replace the sensory receptors of the auditory system and can significantly improve auditory function and quality of life of implanted patients^1^. Likewise, retinal implants aim to replace the function of the sensory receptors of the visual system, the photoreceptors, in retinopathies such as retinitis pigmentosa. While there are different approaches to retinal implants in terms of implantation site and technical realizations (subretinal,^2–4^; epiretinal^5^; suprachoroidal^6^), all use short rectangular pulsatile stimuli to activate the remaining retinal neurons. Patients implanted with a retinal prosthesis regain rudimentary but useful visual functions like the capability to identify, localize and discriminate objects^2, 7^.

Despite this success, there are a number of shortcomings in implant-aided vision, many of which are due to the fact that the electrical stimuli are very different from the physiological signals in the retina, which are much slower and smoother than the pulsatile electrical stimuli^8–12^. Incidental activation of axons passing the stimulation target (e.g. a single retinal ganglion cell (RGC)) severely limits spatial resolution^13–16^, while desensitization of retinal neurons to repetitive electrical stimulation impedes high temporal resolution^17^–19. Acuity of prosthesis-mediated vision therefore reaches only a fraction of healthy visual acuity, leaving patients legally blind despite wearing a retinal implant^20^. Another reason why bionic vision comes short of natural vision is that the image processing mechanisms of the intricate retinal network cannot be triggered selectively by retinal implants; pulsatile stimuli do not separately activate channels like the ON and the OFF pathway^21^.

Different approaches have been suggested to tackle these problems, including sinusoidal stimulation as an alternative to pulsatile stimulation^22^. In wild-type retina, ON and OFF RGCs respond in different phases of low-frequency sinusoidal stimulation; however this phase-preference was assigned to activation of photoreceptors. The clinical relevance of sinusoidal stimulation thus appears to be limited^23^. The use of monophasic half-sinusoidal stimuli of different durations allowed to identify stimulus regimes that maximize the response ratio of ON and OFF RGCs^24–26^.

However, the insight gained by such heuristic searches for optimal stimulus parameters is limited to a particular type of stimulus (i.e. in this case sinusoidal and half-sinusoidal, respectively). One way to more exhaustively look for optimal stimuli for a given sensory modality is white noise analysis^27–29^. This concept has been successfully applied to stimulation with the natural stimulus modality in different sensory systems, such as visual stimulation in the visual system^30^, whisker deflection in the vibrissal system in rats^31^, but also to artificial, electrical stimulation in the visual system^28^. Recently, white noise composed of biphasic electrical current pulses with amplitudes drawn from a Gaussian distribution was used to map the spatiotemporal electrical receptive fields of rat retinal ganglion cells. Subsequently fit linear-nonlinear models accurately predicted RGC responses to electrical stimulation^32, 33^. Linear filters of mouse RGCs could also be recovered with a spatially uniform white noise stimulus consisting of normally distributed subthreshold voltage pulses^34, 35^. The results of these studies demonstrate the feasibility of the white noise analysis approach in electrical stimulation. However, these studies sampled subspaces of the stimulus space, varying pulse amplitude, but not duration or frequency. As white noise analysis can only identify the stimulus that is optimal within the space of provided stimuli, one should try to sample a space that is most likely to contain the true optimal stimulus.

Therefore, we developed and applied a smooth electrical Gaussian white noise current stimulus to cover a space of stimuli that more closely approximate the time scale of physiological signals in the retina. This smooth stimulus simultaneously probes the preferences of RGCs with respect to amplitude, polarity and frequency of an electrical stimulus. Using a high-density CMOS-based microelectrode array for stimulation, we were able to reliably activate RGCs in both wild-type mouse retina (*bl6*) and retina from the *rd10* mouse model of retinal degeneration. We estimated linear filters of cells using two approaches to fitting a linear-nonlinear-Poisson (LNP) model: spike-triggered averaging (STA) and maximum likelihood estimation (MLE). The LNP model accurately predicted RGC responses to electrical stimulation, demonstrating that it captures aspects of retinal processing of electrical stimuli relevant for response generation. The model may be useful for guiding the search for stimuli that improve spatial and temporal resolution of prosthetic-aided vision. The linear filters described here provide a starting point for this search.

## Results

### Simultaneous electrical stimulation and recording using a high-density CMOS-MEA

We used a smooth electrical current stimulus applied by a high-density CMOS-microelectrode-array to stimulate *ex-vivo* preparations of wild-type and photoreceptor-degenerated retina in epiretinal configuration (Fig. 1 a). Our setup allowed us to simultaneously electrically stimulate and record, without the need to remove parts of the recording due to stimulation artefacts. We presented five repetitions of a five second long Gaussian white noise stimulus to flatmount preparations of *wt* and *rd10* retinae. Choosing different configurations of active and silent stimulation electrodes allowed localized stimulation. After the smooth stimulation waveform was removed from the recording, spike-sorting allowed to analyse the retinal ganglion cell responses to the stimulus at the level of individual cells (Fig. 1 b). We evaluated the retinal response in wild-type retina (n = 3, *wt*) and blind retina (n=5, *rd10*) to electrical stimulation by computing the temporal linear electrical filters of the RGCs in a model-based approach.

### Reliable RGC responses to electrical Gaussian white noise stimulation

Retinal ganglion cells in healthy and blind mouse retina responded reliably to stimulation with smooth electrical Gaussian white noise (Fig. 2 b and e). For some cells, firing patterns were nearly identical across trials (Fig. 2 b, third row from below (*wt*); and e, third row from below (*rd10*)); others were more variable in their response (Fig. 2 b, fifth row from below (*wt*); and e, last row (*rd10*)). We quantified the reliability of the RGC response to the electrical stimulus by computing a *reliability index, RI* (see Methods). In *wt* retina, the majority of RGCs (N = 55/87) were entrained to the stimulus with a *RI* larger than a threshold of 0.15 (Fig. 2 d). In *rd10* retina, a smaller percentage of cells were above threshold (N = 28/143, RI>0.15, Fig. 2 g); however, the levels of reliability among above-threshold cells were comparable between *rd10* and *wt* retina.

**Figure 1.**
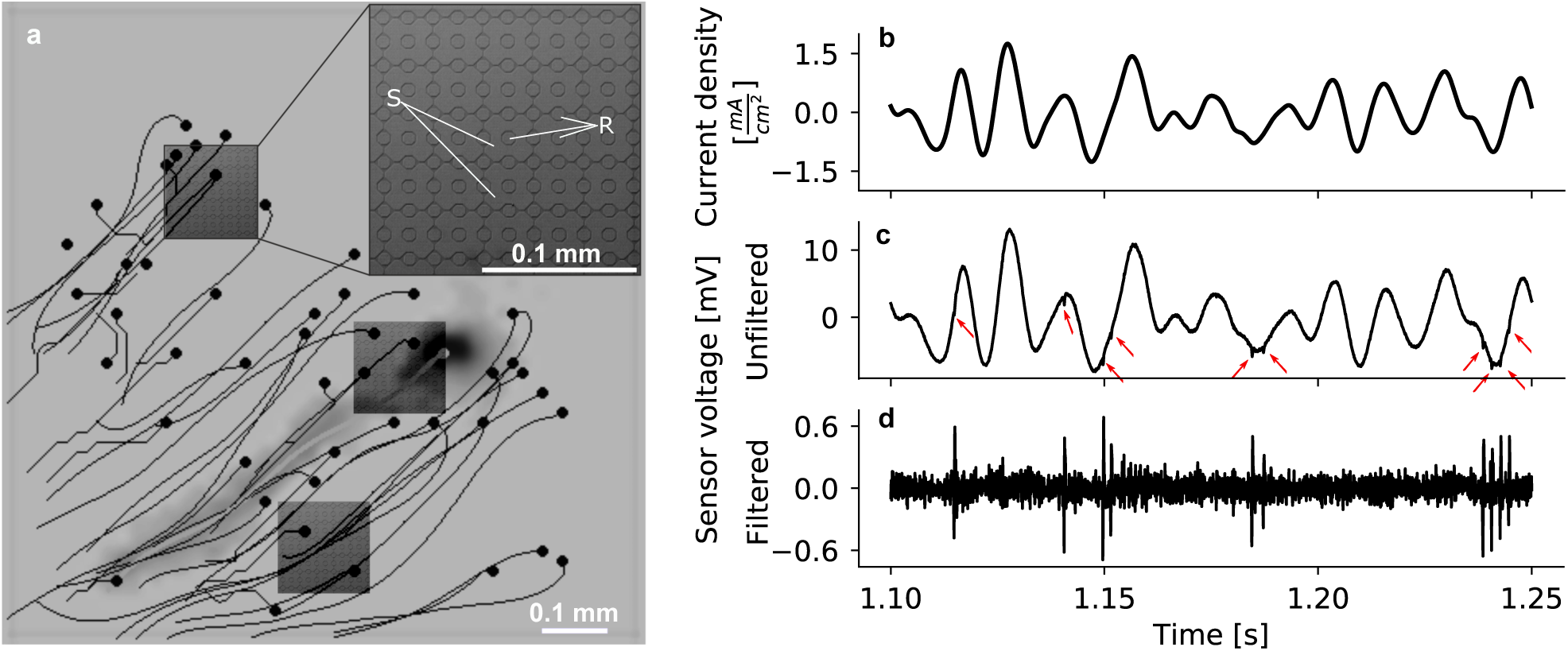
Simultaneous stimulation and recording of RGCs using a hd CMOS-MEA. (a) Schematic of soma location(black dots), axon traces (black lines) of RGCs and configuration of active stimulation areas (dark gray squares) in onerecording. The retina (here from *bl6* mouse) was flat-mounted on a hd CMOS-MEA (gray background). The inset shows thegrid of stimulation electrodes (large elements, labelled S) and recording electrodes (small elements, labelled R). In mostrecordings, only a subset of the stimulation electrodes were active (i.e. delivering the stimulation current). Cell activity wasrecorded simultaneously on recording electrodes. (b) Expected current density of the smooth electrical Gaussian white noisestimulus, calculated as the derivative of the voltage command (see Methods, Eq. 1). (c) Raw recording signal upon stimulationwith the stimulus shown in (b). The stimulus causes an artefact in the raw recording orders of magnitude larger than the signalof interest, the spikes, indicated by red arrows. (d) Signal after filtering with a 2^*nd*^ order band-pass Bessel filter between 1000and 9500 Hz and artefact subtraction. The artefact is removed from the signal and spikes are clearly detectable

**Figure 2.**
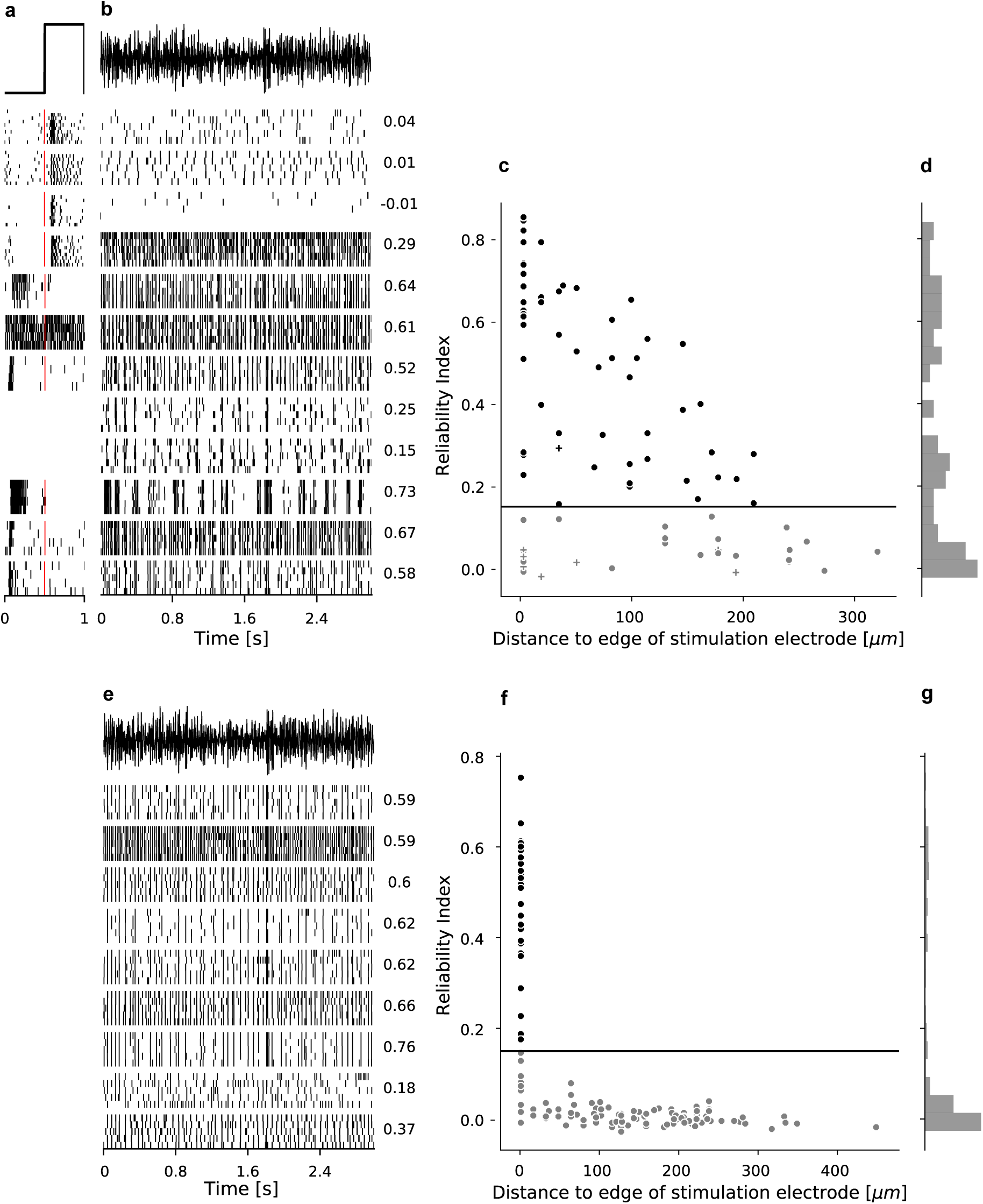
Responses of RGCs to light flashes and smooth electrical stimulation. (a) Raster plot of the responses of RGCs from *wt* mouse retina to a fullfield light flash stimulus. The time course of the stimulus is indicated in the first row, and light onset is marked by a red vertical line in every raster plot. Not every cell was recorded both during electrical and light stimulation. (b) Raster plots of the responses of RGCs from *wt* mouse retina to an excerpt from the smooth electrical Gaussian white noise stimulus (stimulus shown in the first row). Numbers next to each row indicate the reliability index *RI* of the cell’s response to electrical stimulation. (c) Reliability Index as a criterion for reliability of the response is plotted against distance to the edge of the closest active stimulation electrode. Black horizontal line indicates reliability threshold *RI >* 0.15 for inclusion in further analysis. ‘+’ markers indicate cells with ON light response. (d) Histogram of the reliability indices of *wt* RGCs. (e) to (g) sames as (b) to (d) but for RGCs from *rd10* retina.

One major challenge in bionic vision is the selective activation of different information channels such as the ON and the OFF channel^21, 36^. In order to be able to evaluate whether RGCs pertaining to these retinal processing pathways are differently affected by our stimulus, we classified RGCs from healthy retina as ON, OFF or ON-OFF based on their light response. We presented light flashes (0.5 s OFF, 0.5 s ON; see Methods and Fig. 2 a, first row) and recorded the RGC responses to light stimulation. Of the cells that could be identified across recordings with light and electrical stimulation (51/87), 10 cells increased their spiking during incremental light stimuli (ON response, Fig. 2 a, rows 1-4), 17 increased the activity during light decrements (OFF response, Fig. 2 a, row 5, third row from below), and the remaining 24 cells were either ON-OFF cells or had no discernible light response. Only one of the 10 cells with ON response passed the reliability criterion of *RI >* 0.15 during electrical stimulation (see Fig. 2 a and b, row 4), while 16/17 cells with OFF response and 15/24 cells with ON-OFF response responded reliably to electrical stimulation.

The reliability of the RGC response demonstrates that smooth electrical Gaussian white noise can efficiently activate RGCs in healthy and blind mouse retina. The different percentages of activated cells in *wt* and *rd10* retina can be explained by a larger spatial extent of the sensitivity to electrical stimulation in *wt* compared to *rd10* retina. This will be explored in the following paragraph.

### Localized response to local electrical stimulation

Previous studies have investigated the spatial extent of the *electrical receptive field* (spatial eRF), i.e. the area in which electrical stimulation evokes a response in a cell. In healthy rat retinas, eRF with diameters of about 250 *µm* have been reported, while the eRF in blind rat retinas was found to span about 200 *µm* in diameter^4^. Recent studies in healthy and blind mouse retina found eRF diameters of around 400 *µm* for RGCs in healthy mouse, and diameters of around 350 *µm* for RGCs from blind mouse^37, 38^.

We therefore expected the effect of the electrical stimulation on RGC response to be spatially confined. Indeed, we found that the reliability of the RGC response to the electrical stimulation decreased with increasing distance from the closest active stimulation electrode (Fig. 2 c and f). In *wt* retina, the majority of cells within a radius of 200 *µm* of an active stimulation electrode were entrained to the stimulus (*RI >* 0.15). Cells outside this radius did not respond robustly to the stimulus (*RI ≤* 0.15). This is largely in agreement with previously reported eRF sizes. We hypothesized that the small fraction of ON cells responding to electrical stimulation was due to the fact that by chance, the recorded ON cells were located further away from active stimulation electrodes. However, this was not the case; ON cells were recorded at distances from 0 to ≈ 200 *µm* from the closest active stimulation electrode (Fig. 2 c).

In *rd10* retina, cells responded only if they were located directly adjacent to or above an active stimulation electrode (Fig. 2 f). This difference in spatial extent of the sensitivity to electrical stimulation between *wt* and *rd10* suggests that different mechanisms underlie the RGC response to electrical stimulation in healthy compared to blind mouse retina.

### Temporal electrical linear filters of RGCs

We computed temporal electrical linear filters by fitting a linear-nonlinear-Poisson model using two different methods, spike-triggered averaging (STA) and maximum-likelihood-estimation (MLE). We did this for all cells which responded reliably to the stimulus (reliability index *RI >* 0.15, N = 55/87 in *wt* retina, N = 28/143 in *rd10* retina). While filters of RGCs from *rd10* retina were all monophasic negative, filter shapes of RGCs from *wt* retina were more variable (Fig. 3 h,i and m,n).

**Figure 3.**
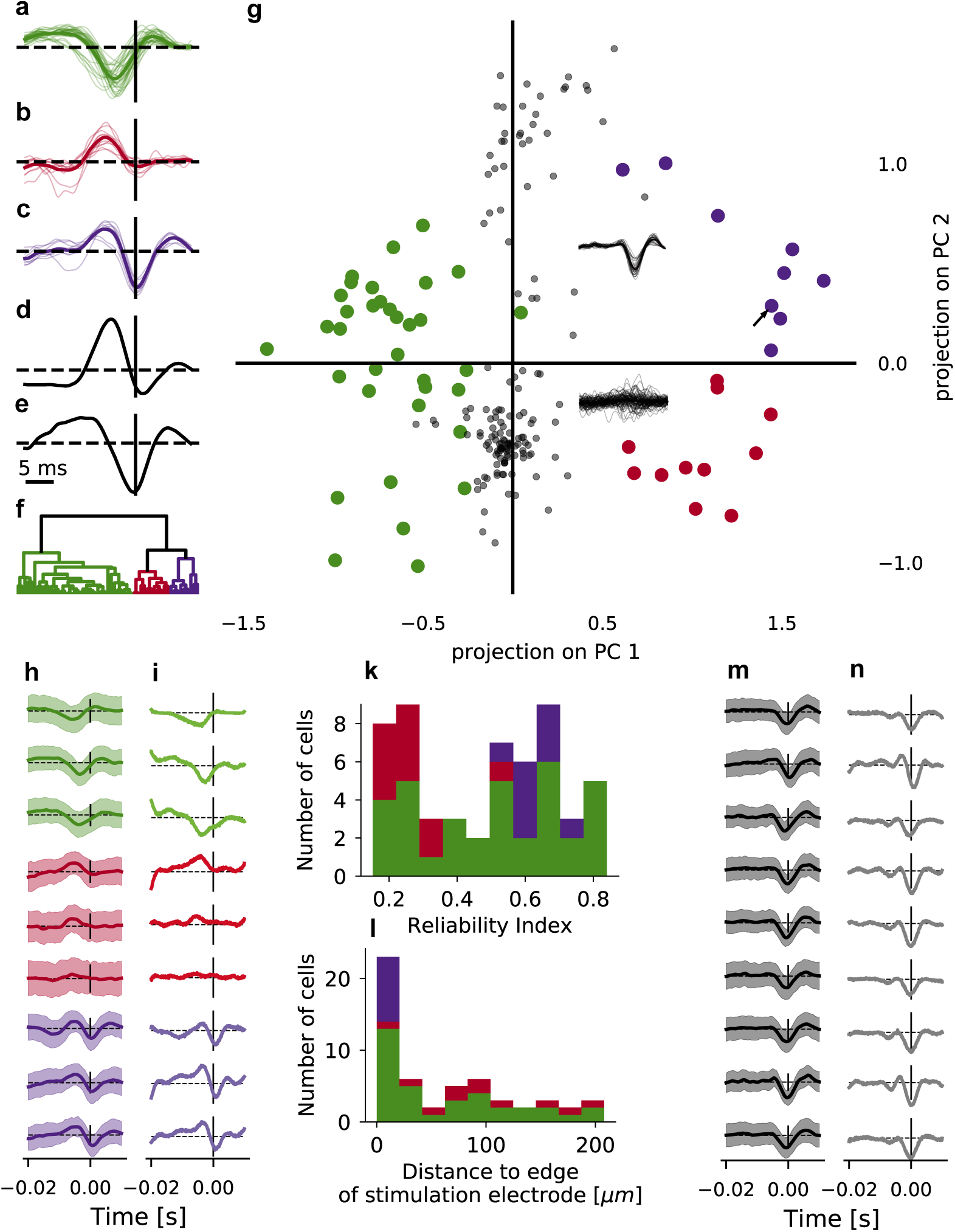
Hierarchical clustering of RGC electrical temporal filters. (a) - (c) Electrical temporal filters of all RGCs from *wt* retina recovered from the STA fit of the LNP model, displayed separately for the three clusters identified by the hierarchical clustering algorithm. Thin lines are individual cell filters, thick lines indicate the average filter for one cluster. (d), (e) 1st and 2nd principal component (PC) recovered from principal component analysis of the ensemble of temporal filters from all cells. (f) Dendrogram showing the separation of consecutively joined clusters along the clustering metric (distance in euclidean space). (g) Scatter plot of the projections of the temporal filters onto the 1st and 2nd PCs (shown in (d) and (e)). Colors indicate cluster assignment. Grey dots indicate the projections of filters of RGCs from *rd10* retina, projected onto the same PCs. The lower inset shows the filters of all *rd10* RGCs whose filter projections are negative in PC2; the upper inset shows the filters of all *rd10* RGCs whose filter projections are positive in PC2. The black arrow marks the cell for which the assignment to clusters did not agree between STA and MLE estimate of the filters. (h) STA estimates of the filters of example RGCs from *wt* retina shown in Fig. 2 b (rows 4-12, same order), with cluster assignment indicated by color. (i) Same as (h), but for MLE estimates of the filters. (k) Stacked histogram of the distribution of reliability indices (RIs) for all cells with *RI >* 0.15, color-coded according to cluster assignment. (l) Distribution of distance to the edge of the closest active stimulation area of all responsive cells (*RI >* 0.15), color-coded according to cluster assignment. (m) STA estimates of the filters of example RGCs from *rd10* retina shown in Fig. 2 e (same order). (n) Same as (m), but for MLE estimates of the filters.

To quantify the difference between the filter shapes found in wild-type retina, we performed a hierarchical clustering on the projection of the filters onto two principal components (PC). For STA filters, the first two PCs (3 d, e) jointly explained 90 % of the variance (PC1: 71.1 %, PC2: 18.9 %). For the MLE filters, the first three PCs explained 62.6 %, 18.4 %, and 11.8 % of the variance; however, we projected onto PC1 and PC3 (Supplementary Fig. S1 d, e), leaving out PC2. We did this because the variance explained by PC2 was mostly due to variance between different recordings, which was not of interest here (see Supplementary Fig. S1 h-j).

We identified two distinct clusters for both STA and MLE filters based on the dendrogram generated by the hierarchical clustering algorithm (see Fig. 3 f and Supplementary Fig. S1 f). These two clusters were separated along the axis of the first PC, and one of them could be split into two sub-clusters that were separated along the axis of the second (STA) or third (MLE) PC. The clusters corresponded to three filter shapes, monophasic negative (green, Fig. 3 and Supplementary Fig. S1 a), monophasic positive (red, Fig. 3 and Supplementary Fig. S1 b) and biphasic (violet, Fig. 3 and Supplementary Fig. S1 c). Assignment to clusters agreed between the STA and the MLE filter estimates for all except one cell. The estimates of the filters of the reliably responding example cells from *wt* retina (Fig. 2 b rows 4-12) fall into these three categories (Fig. 3 h, i). Filters of the example cells from photoreceptor-degenerated retina (Fig. 2 e) were all monophasic negative (3 m, n).

In order to compare the filters of RGCs from *wt* and *rd10* retina, we projected the LNP estimates of the filters from *rd10* RGCs onto the LNP PCs obtained from *wt* retina (Fig. 3 g, gray dots). The densely grouped projections in the lower part of the plot corresponds to projections of flat filters from non-responsive cells, while the more spread-out upper group corresponds to projections of filters from responsive cells (Fig 3 g, insets). Visual comparison of the filter shapes suggests that, while filters from *rd10* RGCs do not fall into one of the three clusters found in *wt*, they are more similar to monophasic negative and biphasic than to monophasic positive *wt* filters (3 h, i and m, n). The pattern of clustering described above confirms this observation. We calculated the latencies of the (positive and/or negative) peaks of the STA and MLE filters of *wt* and *rd10* RGCs relative to the spike (see table 1). In *wt* retina, the latencies of the peaks of monophasic positive filters were around 2 ms longer than the latencies of the peaks of monophasic negative filters. For cells with biphasic filters, the spike occurred within 1 ms of the negative peak of the filter, preceded by 3 to 8 ms by the positive peak. For the monophasic negative filters of RGCs from *rd* retina, the spike occurred within less than 2 ms of the peak of the filter, comparable to the latencies observed for biphasic filters in *wt* retina.

**Table 1.** Peak latencies of filters. The median and the range of the peak latencies relative to spike time for STA filters (top) and MLE filters (bottom). Negative values indicate that the peak occurred before the spike.

|  | latency of neg. peak[ms] | latency of pos. peak [ms] |
| --- | --- | --- |
| monophasic neg. (green, N=35) | -3.7, [-6.1, -1.3]<br>-3.3, [-5.3, -0.6] |  |
| monophasic pos. (red, N=11) |  | -5.5, [-6.2, -4.1]<br>-5, [-6.6, -3.9] |
| biphasic (violet, N=9) | 0.2, [-0.1, 0.7]<br>-0.3, [-0.9, 0.1] | -5.3, [-7.7, -4.5]<br>-3.7, [-7.8, -3.1] |
| *rd10* (N=28) | -0.6, [-1.2, 0.3]<br>-1.2, [-1.8, 0.0] |  |

### LNP model accurately predicts RGC responses to electrical stimulation

We predicted the firing rates of RGCs from *wt* and *rd10* mouse retina using the STA and the MLE fit of the LNP model (see Methods, Fig. 4 a, b (*wt*); Fig. 5 a, b (*rd10*)). The models were fit on 80 % of the data (4 s of the stimulus), and the parameters recovered from this fit were used for the prediction of the remaining 20 % of the data (1 s of stimulus). For RGCs from *wt* retina, prediction performance *P*^*wt*^, evaluated as the correlation between true and predicted firing rate (Methods, Eq. 22) ranged from 0.05 to 0.67 for the STA fit and from 0.1 to 0.7 for the MLE fit of the model (Fig. 4 c, d). The MLE fit performed slightly better than the STA fit (mean 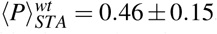, mean 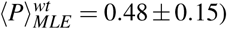, and both models performed better for cells with monophasic negative and biphasic filter shapes compared to cells with monophasic positive filter shapes (Table 2 and Fig. 4 e).

**Table 2.** Prediction performance of the LNP model (STA and MLE fit) Prediction performance (mean ±s.d., [range]) of the STA fit and the MLE fit, measured as the correlation coefficient 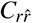 (see Methods) between predicted and true firing rate at a resolution of 1 kHz, evaluated separately for the three identified clusters among *wt* filters, as well as for *rd10* filters.

|  | STA fit performance | MLE fit performance |
| --- | --- | --- |
| negative-only (green, N=35) | $0.52 \pm 0.09$, [0.32, 0.67] | $0.54 \pm 0.09$, [0.35, 0.70] |
| positive-only (red, N=11) | $0.21 \pm 0.11$, [0.05, 0.41] | $0.25 \pm 0.11$, [0.10, 0.42] |
| biphasic (violet, N=9) | $0.49 \pm 0.06$, [0.39, 0.58] | $0.52 \pm 0.06$, [0.42, 0.62] |
| *wt* all (N=55) | $0.46 \pm 0.15$, [0.05, 0.67] | $0.48 \pm 0.15$, [0.10, 0.70] |
| *rd10* all (N=28) | $0.53 \pm 0.08$, [0.35, 0.66] | $0.55 \pm 0.1$, [0.37, 0.69] |

**Figure 4.**
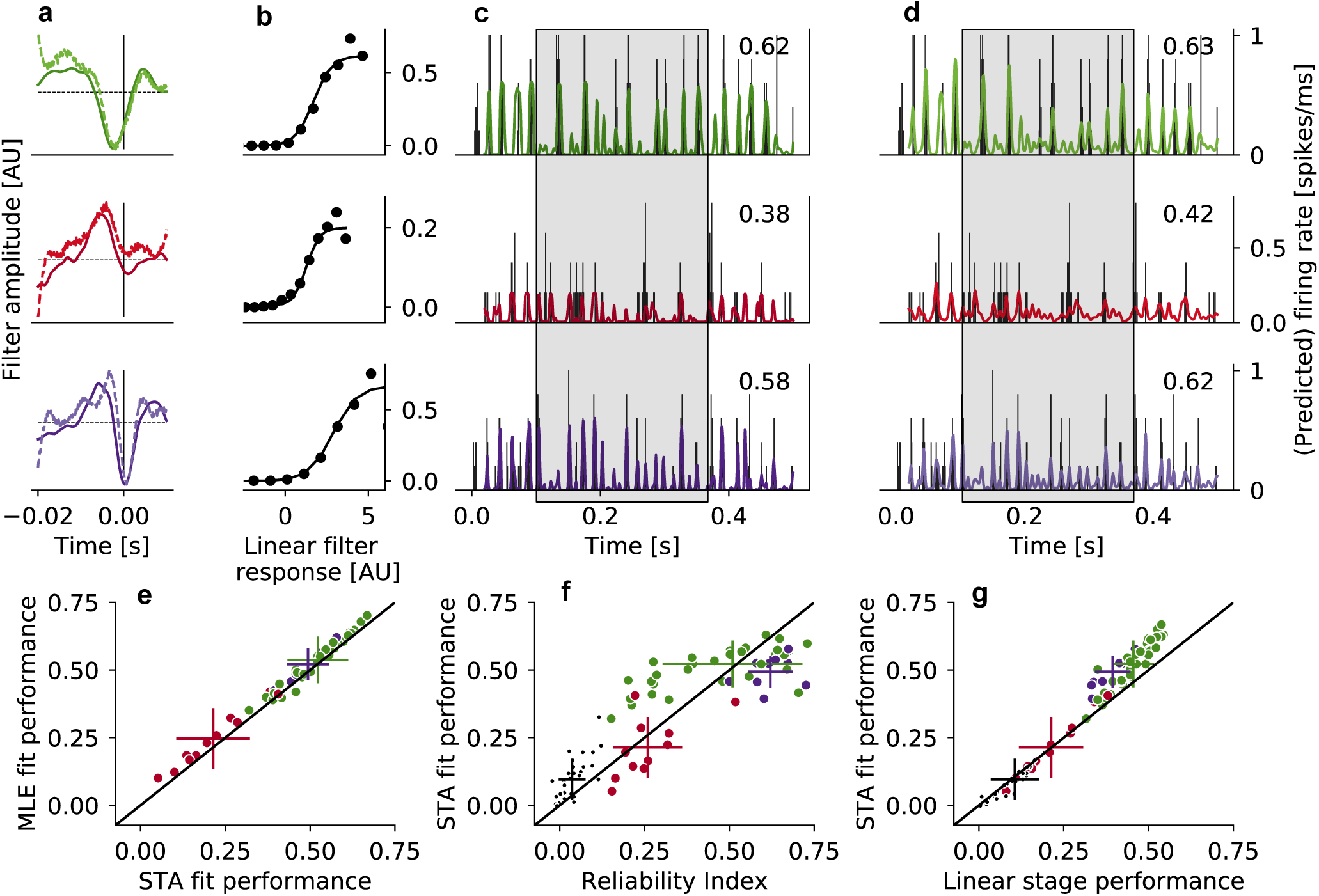
Model prediction of firing rates in *wt* retina. (a) Linear filters of three example cells from different clusters, derived from the STA fit (solid lines) and from the MLE fit (dashed lines); displayed at different scales in arbitrary units. To create the firing rate prediction, the filter response of the stimulus snippets with these filters was computed, and then the nonlinearity was applied. (b) Nonlinearity of the STA model; the nonlinearity of the MLE was the standard sigmoid function. (c) True firing rate, binned into 1 ms bins (black histogram) and the firing rate prediction of the STA fit of the model (colored traces) for three example cells; the highlighted region shows differential responses of cells with different types of filters, which are correctly predicted by the model. Decimal numbers in upper right corner indicate the performance of the model for that specific cell. (d) Same as (c) for the MLE fit of the model. (e) Prediction performance of the MLE fit plotted against the prediction performance of the STA fit; colors indicate cluster assignment. (f) Prediction performance of the STA fit plotted against reliability index. Colors indicate clusters assignment. Small black dots represent RGCs that did not pass the reliability criterion of *RI >* 0.15, but are shown here for completeness. (g) Prediction performance of the STA fit of the full LNP model plotted against the prediction performance of only the linear stage of the LNP model. Colors indicate cluster assignment. Small black dots represent RGCs that did not pass the reliability criterion of *RI >* 0.15, but are shown here for completeness.

**Figure 5.**
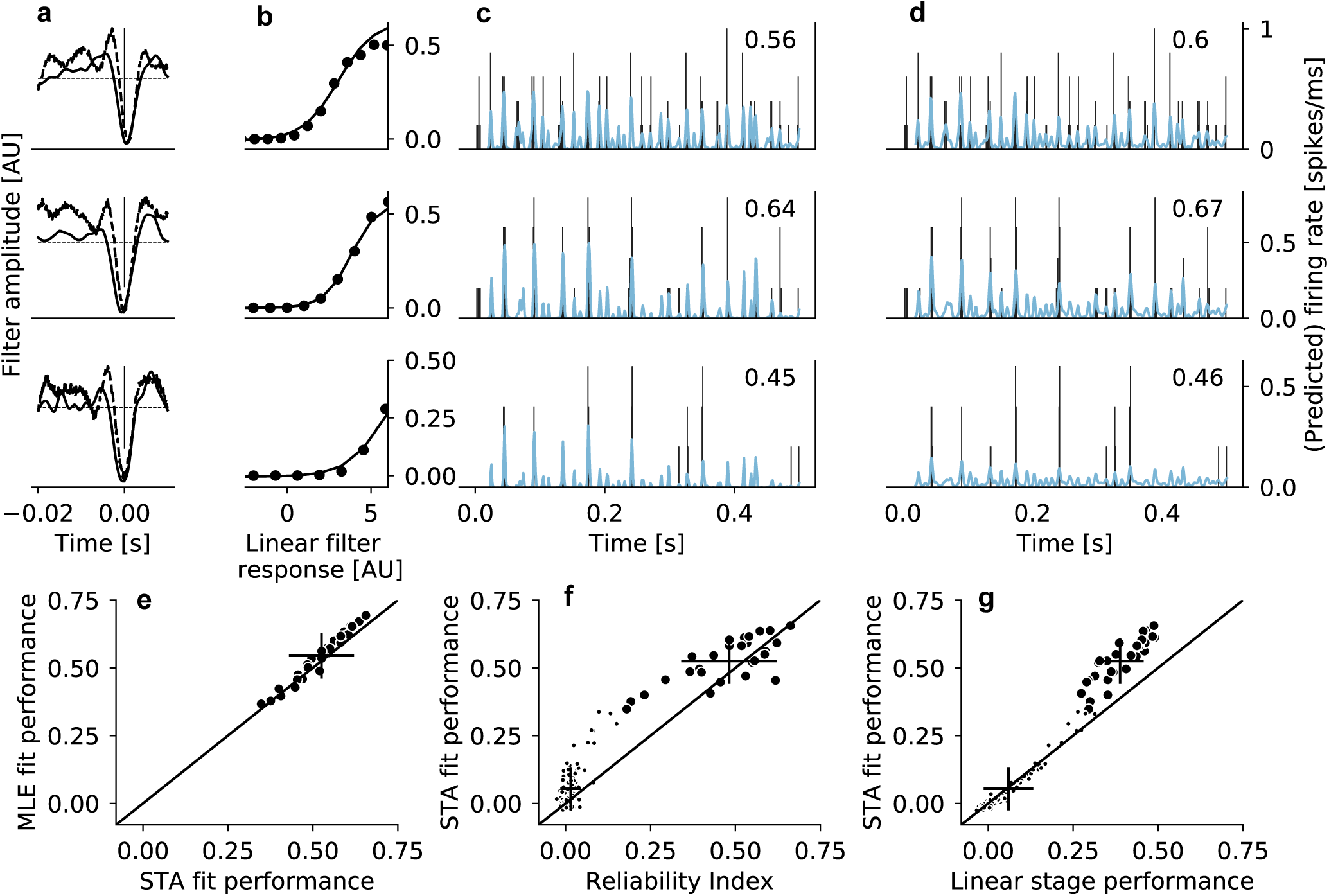
Model prediction of firing rates in *rd10* retina. (a) Linear filters of three example cells, derived from the STA fit (solid lines) and from the MLE fit (dashed lines); displayed at different scales in arbitrary units. To create the firing rate prediction, the filter response of the stimulus snippets with these filters was computed, and then the nonlinearity was applied. (b) Nonlinearity of the STA model; the nonlinearity of the MLE was the standard sigmoid function. (c) True firing rate, binned into 1 ms bins (black histogram) and the firing rate prediction of the STA fit of the model (blue traces) for three example cells. Decimal numbers in upper right corner indicate the performance of the models for that specific cell. (d) Same as (c) for the MLE fit of the model. (e) Prediction performance of the MLE fit plotted against the prediction performance of the STA fit. (f) Prediction performance of the STA fit plotted against reliability index. Small black dots represent RGCs that did not pass the reliability criterion of *RI >* 0.15, but are shown here for completeness. (g) Prediction performance of the STA fit of the full LNP model plotted against the prediction performance of only the linear stage of the LNP model. Small black dots represent RGCs that did not pass the reliability criterion of *RI >* 0.15, but are shown here for completeness.

We hypothesized that there was a positive relationship between the prediction performance of the LNP model and the reliability of a cell’s response, indicated by its reliability index *RI*. Indeed, the performance of the model was generally better for more reliable cells (Fig. 4 f). However, there were differences in model performance between cells with different filter shapes that could not be explained by differences in their response reliability; model performance was worse for cells with monophasic positive filters than for cells with monophasic negative filters, even when the cells were equally reliable in their response. Thus, the difference in model performance must either be due to differences in the quality of fit of the linear stage, the nonlinear stage, or a combination of both. To disentangle the effects of the linear and the nonlinear stage on model performance, we compared the accuracy of only the linear stage to that of the full model in predicting the firing rate. The linear stage of the LNP model can be used to predict the firing rate by computing the dot product between the linear filter and the stimulus, without applying the static nonlinearity. Differences in performance between RGCs with monophasic negative and biphasic filters on the one hand and RGCs with monophasic positive filters on the other hand already emerge at the linear stage (Fig. 4 g, x-dimension). The prediction performance further diverges when applying the nonlinearity (Fig. 4 g, y-dimension). Thus, differences in model performance between cells are a cumulative effect of differences in the reliability of the cells’ response, but also of differences in the quality of the fit of the linear stage as well as the nonlinear stage.

Different filter shapes are the result of different response patterns that are elicited by the stimulus. Interestingly, both model fits also predict differences in response patterns between cells with different filter shapes (Fig. 4, c and d, highlighted region).

As noted before, linear filters of RGCs from *rd10* retina were most comparable to monophasic negative and biphasic filters of RGCs from *wt* retina (Fig. 3 and Fig. 5 a). Likewise, patterns of model performance were similar in *rd10* RGCs and *wt* RGCs with monophasic negative and biphasic filters. For RGCs from photoreceptor-degenerated retina, the performance of the STA fit of the model ranged from 0.35 to 0.66 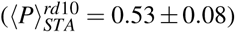; again, the MLE fit performed slightly better (range 0.37 to 0.69, 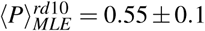, Fig. 5 c, d and e). These performance values are very similar to those obtained for RGCs with monophasic negative filters from *wt* retina. Furthermore, the relationship between reliability of the response and LNP performance (Fig. 5 f), as well as the relationship between linear stage performance and full model performance (Fig. 5 g) in *rd10* retina were similar to these relationships for RGCs with monophasic negative and biphasic filters from *wt* retina (Fig. 4 f, g).

## Discussion

Retinal implants represent one of the most promising treatment options for patients suffering from degenerative retinal diseases like retinitis pigmentosa or macular degeneration, but clinical efficacy remains limited due to non-physiological and therefore suboptimal stimulation strategies. The goal of this study was to investigate the response properties of retinal ganglion cells to a novel electrical stimulus that comes closer to the physiological signals in the retina. Identifying an electrical stimulus that is both physiologically plausible and technically feasible might significantly improve clinical outcomes in patients wearing retinal implants.

We electrically stimulated retinal ganglion cells from wild-type and photoreceptor-degenerated mouse retina with smooth Gaussian white noise currents and used a model-based approach to estimate linear filters of the RGCs. The estimates of the linear filters could be clustered into three different groups based to their shapes and constituted the linear stage of a linear-nonlinear-Poisson model, which accurately predicted retinal ganglion cell firing probability. This study demonstrates that physiologically plausible electrical stimuli can be used to activate retinal ganglion cells in both wild-type and photoreceptor-degenerated retina; furthermore, the LNP model could be used to find stimuli that maximize the response in cells with one class of filters while minimizing the response in cells with a different class of filter.

One of the main goals in retinal prosthetics is the selective activation of different information channels in the retina, most prominently the ON and OFF channels. A prerequisite for achieving this goal are cell-type specific preferred electrical stimuli, like it is the case for light stimuliation of healthy RGCs^36, 39^. Indeed, investigating network-mediated electrical activation of rat RGC, ON cells were found to prefer cathodic-first biphasic pulses, and OFF cells to prefer anodic-first pulses, but the difference in preference was small^33^. Antagonistic polarities of electrical filters in wild-type mouse ON and OFF cells have been reported as well^35^; however, the clinical applicability of these filters as stimuli is limited as they most likely arose due to photoreceptor activation^40^.

In this study, we found three different classes of RGC filters in response to smooth electrical stimulation in wild-type, and one class of filter in photoreceptor-degenerated mouse retina. As only one ON cell responded reliably to the electrical stimulus, we can not draw conclusions about different optimal stimuli for ON and OFF pathway activation; more data from ON cells needs to be collected. However, the fact that we find different classes of filters even within one information channel, that is, the OFF channel, suggests that much more fine-grained selective activation of RGC types, for example sustained or transient types, may be possible with the use of physiologically plausible stimuli derived from the filters presented here.

We found different classes of filters in healthy but not in blind retina, which might cast doubt on the clinical relevance of our findings. Specifically, if our electrical stimulus activated RGCs in healthy retina via photoreceptors, and the different filter shapes were due to the sign-preserving and sign-inverting synapses between photoreceptors and OFF and ON bipolar cells, respectively^35, 40^, selective activation of the ON and OFF pathways would not be achievable in blind retina. Therefore, identifying which retinal elements (photoreceptors, bipolar cells or RGCs) are activated by our stimulus is highly relevant. One indicator of the origin of RGC responses to electrical stimulation is the latency of spikes relative to the stimulus. As our stimulus is time-continuous, we cannot directly compute the response latency as the time difference between stimulus delivery and response. Instead, we used the latencies of the filter peaks relative to the spike time as a proxy for response latency. The latencies of the filter peaks (see Table 1) are much shorter than the latencies > 40 *ms* usually reported for photoreceptor-mediated activation of ganglion cells in rabbit and rat retina^24, 41, 42^. It is therefore unlikely that the RGC responses in *wt* retina were due to activation of photoreceptors. Rather, the latencies of both positive and negative monophasic filter peaks fall in the range usually reported for bipolar cell activation^38, 43^, whereas the short latencies of the negative peak of biphasic filters suggest that these filters arose due to direct ganglion cell activation^16, 44^. Only cells within or directly adjacent to an active stimulation area showed biphasic filters, while cells with negative or positive monophasic filters responded within a radius of *≈* 200 µ *m* of stimulation electrodes, congruent with reports of the typical size of electrical receptive fields in mouse retina^37, 38^. Taken together, the properties of biphasic filters suggest that these were the result of direct retinal ganglion cell activation, while both monophasic filter shapes were more likely to arise due to bipolar cell activation.

RGC responses from photoreceptor-degenerated retina were most comparable to the RGCs from wild-type retina with biphasic filters in terms of latency and shape of the filters, and spatial extent of the sensitivity to electrical stimulation (Table 1, Figs. 2, 3). It is therefore conceivable that different retinal elements are activated in wild-type and photoreceptor-degenerated retina by smooth electrical stimulation at the given intensity: bipolar and ganglion cells in wild-type retina, and only ganglion cells in photoreceptor-degenerated retina. As the retinal network initially remains intact in the *rd10* model of retinal degeneration as well as in patients suffering from retinitis pigmentosa^37, 45–47^, it should be investigated whether bipolar cell-mediated RGC activation can be achieved in photoreceptor-degenerated retina with our stimulus. Experiments with retina from younger (and therefore less degenerated) *rd*10 mice, as well as stronger stimulation intensities to account for potentially higher thresholds in *rd* models^48, 49^ will reveal whether different filter classes can be recovered in blind mouse retina.

Our stimulus could prove clinically relevant even if different filter classes cannot be identified in RGC from *rd10* retina. Responsive neurons are confined to locations directly adjacent to or within the stimulation area (Fig. 2 f). Furthermore, the latencies of the filter peaks in this study were much shorter than the peak latencies found in previous studies using different stimulation protocols. Epiretinal stimulation of wild-type mouse retina with subthreshold pulsatile Gaussian white noise yielded biphasic filters with peak latencies roughly 2 orders of magnitude longer than the filters described here^34, 35^. Filters estimated from network-mediated rat RGC responses to photovoltaic stimulation were about 10 times slower^50^. The short temporal scale of the filters estimated from RGC responses to smooth Gaussian white noise is a result of fast RGC responses to electrical stimulation. A stimulus derived from the filters described here, implemented in a retinal prosthesis, might therefore grant precise control over temporal and spatial activation patterns and thus help to overcome problems such as elongated phosphenes^51^ and fading^18^.

Attempts of modelling a neural system of interest are ubiquitous in neuroscience in general and in neural prosthetics in particular. An appropriate encoding model allows to predict the system’s response to an input, and to draw inferences about the elements of the system^52–54^. Furthermore, it can be used to find optimal stimuli that maximally activate a target neuron^55^. Our model predicts differences in electrically evoked responses of cells with different filter shapes (see Fig. 4, c,d, highlighted region) and might therefore be used to identify stimuli that will maximize the response of cells with one type of filter while minimizing the response in cells with a different type of filter. Identifying such stimuli might potentially provide a tool for selective activation of certain cell types. However, as different filter shapes could not be associated with defined RGC types such as ON or OFF RGC types, and as different filter shapes were so far only identified in wild-type retina, future work will reveal to what extent this approach will provide cell-type specific stimulation protocols for retinal prostheses.

## Methods

All experimental procedures were carried out in compliance with §4 of the German law on animal protection and were approved by the Regierungspräsidium Tübingen (Registration No.: 35/9185.82-7). All the experiments were performed in accordance with the ARVO statement for the use of animals in ophthalmic and visual research.

### Retina preparation

*Ex vivo* retina from five B6.CXB1-Pde6brd10/J (*rd10*) mice (2 female, 3 male; age between p80 and p209) and from three C57BL/6J (*wt*) mice (2 female, 1 male; age between p87 and p274) was used. Animals were dark-adapted for approximately 30 minutes before the experiment, anaesthetised with carbondioxide and euthanized by cervical dislocation. Both eyes were then enucleated, the retina was isolated and dissected in Ames’ buffer under dim red-light conditions to prevent bleaching of remaining photoreceptors (further details have been described previously^37, 56^). A retinal portion of ca. 1-2 *mm*^2^ was placed on a CMOS-based microelectrode array in epiretinal flat-mount configuration. Before each use, the microchip was cleaned with Terg-a-zyme (Sigma Aldrich, Z273287 dissolved in bidistilled water) and then coated with Poly-L-lysine (Sigma Aldrich, P2636, 1 mg/ml dissolved in bidistilled water). The recording chamber (2 *ml*) was perfused with warm, carbonated Ames’ buffer (Sigma Aldrich, A1420, 36°C, pH 7.4) at a flow rate of 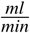

### High-density CMOS-based microelectrode array (hd CMOS-MEA)

Stimulation and recording of retinal ganglion cell activity were performed with a high-density CMOS-based microelectrode array (CMOS MEA 5000, Multi Channel Systems MCS GmbH, Reutlingen, Germany). The MEA comprised 1024 capacitive stimulation electrodes and 4225 recording electrodes on an area of 1 *mm*^2^ (see Fig. 1). Each stimulation electrode extended over an area of 688 *µm*^2^ with 0.5 *µm* spacing between stimulation electrodes. The stimulation electrodes were made of a titanium nitrite and were covered by a native oxide layer which insulated the metal from the electrolyte in the recording chamber. The pitch (center-to-center spacing) between two recording electrodes was 16 *µm*. Details of the CMOS-based MEA have been reported previously^57, 58^.

### Electrical stimulation

We designed an electrical stimulus consisting of Gaussian white noise low-pass filtered at 100 Hz with a 5^*th*^ order Butterworth filter (Fig. 1 b). The electrical stimulus was applied using the hd CMOS-MEA 5000 (see Fig. 1 a). Applying voltages to the stimulation electrodes of the chip evokes capacitive currents in the electrolyte. The stimulation current density is proportional to the time derivative of the electrode voltage and scales with the specific electrode capacitance *c*:

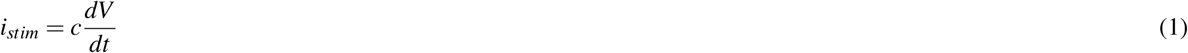

Therefore, in order to obtain a certain current *i*_*stim*_, the integral function of the desired current has to be applied to the chip as voltage command. For our electrical Gaussian white noise stimulus, this was achieved by generating *f*_*s*_ *T* samples (where *f*_*s*_ is the sampling frequency and *T* is the total stimulus duration) from a standard normal distribution, and then calculating the cumulative sum over these samples. To reduce the drift in the random walk that is generated by this process, a reflection limit *L*_*reflect*_ was introduced: whenever adding a new sample to the cumulative sum would raise the absolute value of the cumulative sum above the reflection limit *L*_*reflect*_ = 10, the step’s sign was inverted. This yielded a sequence of values (in arbitrary units) describing a random walk, which was then rescaled to generate a voltage command *V*_*command*_ that respects the safe input limits of the stimulus generator, [0*V ≤ V*_*command*_*≤* 2.5*V*]. The introduction of the reflection limit reduces the power of the stimulus in the low frequency regime; apart from that, power spectral density of the resulting signal is relatively flat up to a frequency of 100 Hz and then drops off (Fig. 6 b; note that frequencies < 4 Hz are not shown to improve visibility).

**Figure 6.**
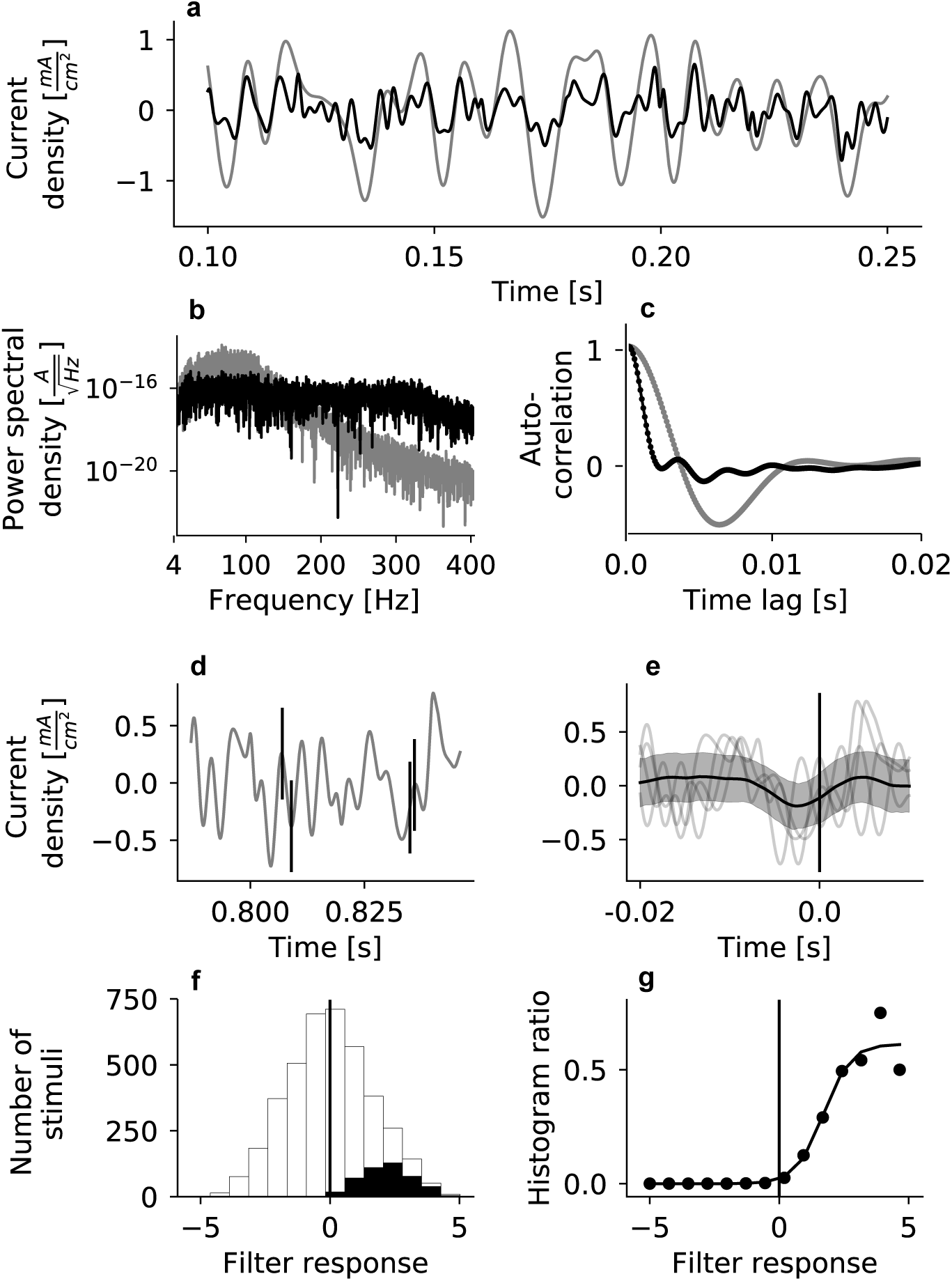
LNP model: Stimulus whitening and filter estimation by spike-triggered averaging. (a) Example segment of the original stimulus in grey, and the corresponding segment of the whitened stimulus in black. (b) Power spectral density (PSD) of the original (grey) and whitened (black) stimulus. While the PSD of the original stimulus declines at frequencies > 100 Hz due to filtering, this effect is undone by whitening. (c) The autocorrelation function of the original (grey) and the whitened (black) stimulus at time lags of up to 20 ms. The temporal extent of the autocorrelation of the whitened stimulus is much reduced compared to the original. (d) For every spike of a RGC (black vertical lines), a segment of the whitened version of the stimulus, spanning 20 ms before and 10 ms after the spike, is added to the spike-triggered ensemble (STE). (e) The spike-triggered average (black trace) is computed by averaging across all elements of the STE (the STE elements from panel (a) are shown as thin gray lines). Gray shaded area indicates *±* 1 standard deviation of the STE. Vertical black line indicates time of spike.(f) The elements of the full stimulus ensemble, consisting of all 300 sample long stimulus snippets taken 10 samples apart, are projected onto the linear filter of a cell and binned to yield a histogram (empty bars). The same is done for the elements of the STE (black histogram). (g) The nonlinearity (black dots) is estimated as the ratio between the histogram of the projected elements of the spike-triggered ensemble to the histogram of the projected elements of the full stimulus ensemble (black and the empty histograms from panel (f), respectively) and fit by a sigmoidal (Eq. 12) or exponential (Eq. 13) function (black trace, here sigmoidal).

The stimulus applied in the experiments described here was generated using the parameters *f*_*s*_ = 10 kHz, *T* = 5 *s* and *L*_*reflect*_ = 10 (*AU*). The current density was estimated as described before^38, 41^. The maximal current density achieved with these parameters and the available chips was 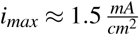.

### Light stimulation

To probe the light response of RGCs in wild-type retina, we presented full-field light flashes repeated 4 to 10 times. Light stimuli were generated by a selected LED (pE 4000 coolLED, peak wavelength 470-490 *nm*) and were flashed on for 500 ms, then turned off for 500 ms, thus creating a 1 Hz flicker. The light stimuli were projected onto a custom-made digital mirror display (*µ*-Matrix, Rapp Optoelectronic GmbH) mounted on an upright microscope (Olympus, BX51WI). The digital mirror display was focused onto the backside of the microscope objective.

### Artefact reduction and preprocessing

In the raw recording, spiking activity is masked by large stimulation artefacts, making spike sorting and further analysis impossible (Fig 1 b). Therefore, a series of data preprocessing steps was performed to remove the stimulation artefact. First, the raw traces of each recording electrode were band-pass filtered between 1000 and 9500 Hz with a 2^*nd*^ order Bessel filter. Then, the contribution of the stimulation artefact to the raw trace was estimated by computing the dot product between the applied stimulus *S* and the raw trace *r*_*raw*_ (*r*_*raw*_, *S* are N-dimensional column vectors), normalized to the dot product of the stimulation waveform with itself:

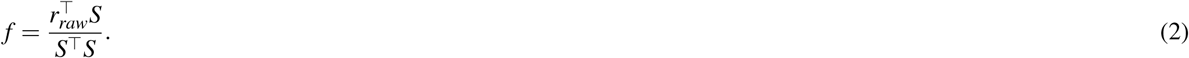

To remove the contribution of the stimulation artefact in the raw recording, the stimulus trace (Fig. 1 b) was then subtracted from the raw recording (Fig. 1 c), multiplied by the factor *f*:

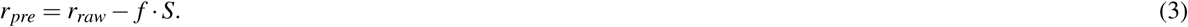

In the resulting signal, spikes are clearly separated from the background signal and no longer masked by the stimulation artefact (Fig. 1 d). Spike sorting was performed on this signal using the spike sorter based on convolutive independent component analysis^59^, implemented in the MultiChannel Systems CMOS-MEA-Tools Software (https://www.multichannelsystems.com/software/cmos-mea-tools, version 2.1.0).

### Evaluating reliability of RGC responses to electrical stimulation

In order to quantify how reliably RGCs respond across repetitions of the same stimulus, we computed a *Reliability Index* (*RI*). To compute the *RI*, individual RGC responses from 4 repetitions were binned into 2 *ms* wide bins, resulting in a histogram *H*. The same was done for the remaining 5^*th*^ repetition, yielding a histogram *h*. Then the correlation coefficient between the two histograms was computed. This was repeated for all possible combinations of *H* and *h*, and the *RI* was computed as the average across the resulting correlation coefficients:

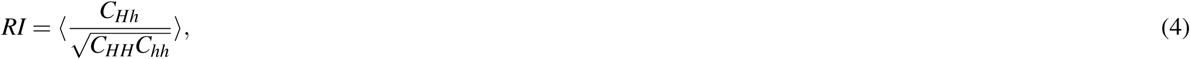

where *C* denotes the covariance matrix of the histogram *H* of cell responses during four repetitions of the stimulus and the histogram *h* of the cell responses during the fifth repetition, and angular brackets ⟨ ⟩ denote the mean across all (five) possible combinations of *H* and *h*. A value of 1 indicates perfect reliability (identical response pattern in all repetitions), while a value close to 0 indicates no reliability in the response to the stimulus. Based on visual inspection of the distribution of the RIs of wild-type RGCs, we included RGCs in the analysis if their RI was larger than 0.15 (see Fig. 2 d). The same threshold was applied to RGCs from photoreceptor-degenerated retina (see Fig. 2 g).

### Fitting a linear-nonlinear-Poisson model to electrically evoked RGC responses

We modeled RGC responses to electrical stimulation with a Linear-nonlinear-Poisson (LNP) model^30^. The LNP model assumes that the firing rate of a neuron in response to a stimulus *s*_*t*_ at time *t* can be modelled as an inhomogeneous Poisson process with instantaneous firing rate

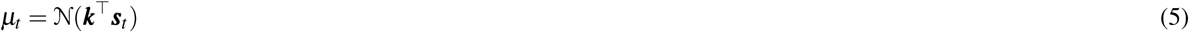

where ***k***^*T*^ ***s***_*t*_ denotes the dot product of the cells linear filter and the stimulus at time t, and 𝒩 denotes a static nonlinearity.

We used two different approaches to fit the LNP, spike-triggered averaging with subsequent estimation of the nonlinearity, and maximum-likelihood estimation of a generalized linear model (GLM) with logit link function.

### LNP model: Filter estimation by spike-triggered averaging

The spike-triggered average^29, 60^ (STA) is calculated by averaging over the spike-triggered stimulus ensemble (STE), which is the collection of all stimuli that were followed by a spike within a defined time window (here 1 *ms*). For a discrete time shift *τ*_*i*_ before the spike the STA is given as

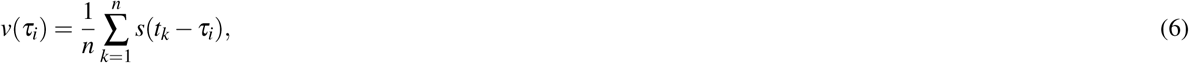

with

*n* the number of spikes,

*s*(*t*) the value of the stimulus at time *t*,

*τ*_*i*_ the time shift relative to the spike.

A cell’s linear temporal filter is then defined as the vector

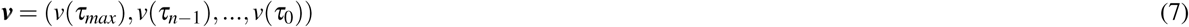

The spike-triggered average was calculated 20 ms into the past and 10 ms into the “future” relative to the spike, i.e. *τ*_*max*_ = –20 ms and *τ*_0_ = 10 ms. The duration of the filters presented here is thus 30 ms, equivalent to 300 stimulus samples.

The stimulus was low-pass filtered at 100 Hz (see Methods) and therefore has a non-zero autocorrelation at time shifts *≤* 10 *ms* (see Fig. 6 c). These autocorrelations affect the shape of the filters computed by spike-triggered averaging^61^. To reduce the effect of the autocorrelations on the filters, the STA was calculated using whitened stimulus “snippets“. For the whitening procedure^62^, the 50 000 samples (5 s) long continuous stimulus was sliced into snippets of 300 stimulus samples, taken 10 samples apart (1 ms). This yielded a stimulus ensemble containing 4701 stimulus snippets, and the covariance matrix was computed based on this ensemble. The square-root of the pseudo-inverse of the resulting covariance matrix, 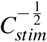, was computed, and each stimulus snippet was whitened by multiplying with 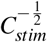:

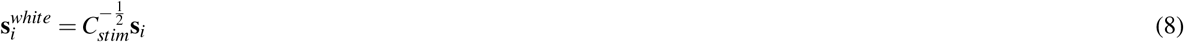

with

**s**_*i*_ *∈ ℝ* ^300*x*1^ stimulus “snippets” of 300 frames each

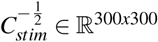 the pseudoinverse of the stimulus covariance matrix *C*_*stim*_.

The pseudoinverse is obtained from the eigendecomposition of the stimulus covariance matrix

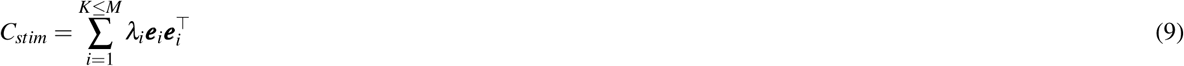

by keeping the eigenvectors ***e***_*i*_*∈ ℝ*^300*x*1^, but inverting and taking the square root of the *L < K* largest eigenvalues *λ*_*i*_ and setting the remaining eigenvalues to zero:

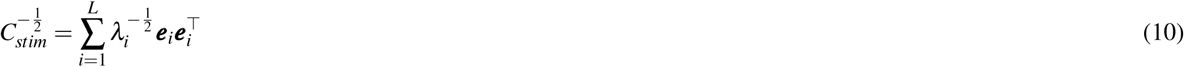

The whitened stimulus used for spike-triggered averaging (Fig. 6 a and d) was reconstructed by concatenating the resulting whitened stimulus snippets and smoothing the edges with a hanning window.

The nonlinearity ℱ was estimated as the ratio between the histograms of the projection of the STE onto the cells linear filter and the projection of the raw stimulus ensemble onto the cells linear filter^30^ (Fig. 6, f and g):

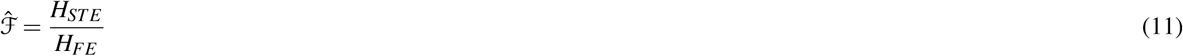

*H*_*STE*_ denotes the histogram of projected stimulus snippets from the spike-triggered ensemble (STE) and *H*_*FE*_ denotes the histogram of projected stimulus snippets from the full ensemble (FE). The estimate of the nonlinearity was then fit by a sigmoidal or exponential function. The sigmoidal fit to the nonlinearity at time *t* is described by the equation

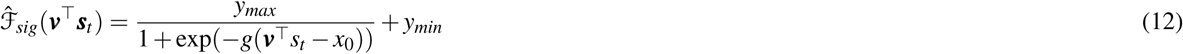

with free parameters *y*_*max*_ (saturation amplitude of the sigmoidal curve), *g* (gain factor), *x*_0_ (50 % threshold of the saturation amplitude) and *y*_*min*_ (vertical offset). The exponential fit to the nonlinearity at time *t* is described by the equation

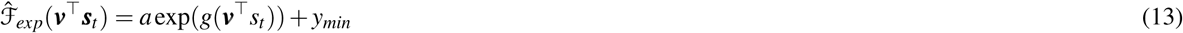

with free parameters *a* (controlling the height of the curve), *g* (gain factor), and *y*_*min*_ (vertical offset). *s*_*t*_ is a whitened stimulus snippet as defined in Eq. 8, containing the 300 stimulus samples preceding *t*.

### LNP model: Filter estimation by maximum-likelihood estimation

In addition to the spike-triggered approach, the temporal electrical filter was also estimated by fitting a GLM with logit link function and an elastic net penalty to the data^63, 64^. Here, the filter coefficients are found by minimizing the negative log-likelihood of the observed data with respect to the coefficients. To constrain the coefficients and to prevent overfitting, a penalty term is added to the loss function. The GLM was fit on the raw (not whitened) stimulus, because the process of maximum likelihood estimation intrinsically performs correlation-correction on the filter estimate^65^. We used a slightly modified version of the `pyglmnet` python package (available at https://github.com/glm-tools/pyglmnet) for fitting the GLM, and a slightly modified version of the `sklearn` python package for cross-validation.

The firing probability *µ*_*t*_ at time *t* is assumed to be related to the stimulus by a sigmoid function

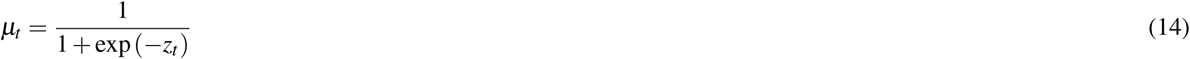

where

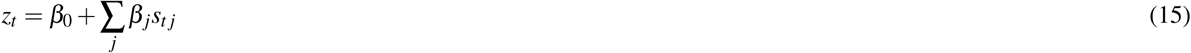

with *j*

*β*_0_ ∈ ℝ a scalar offset

***β*** ∈ ℝ^*m*^ the estimate of the m-dimensional linear filter

is a linear transformation of the stimulus ***s***_*t*_ ∈ ℝ^300*x*1^ at time *t*. Assuming that a spike train *Y* = {*y*_1_, *y*_2_, …, *y*_*t*_}, *y*_*i*_ ∈ 0, 1 of one particular neuron is generated by a sequence of independent (but not identically distributed) Bernoulli trials with time-dependent firing probability *µ*_*t*_, the probability of the spike train is given by

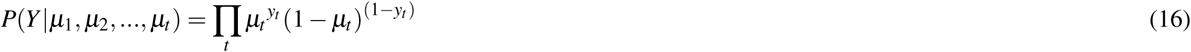

The average log-likelihood function ℒ (***µ***;*Y*) of the parameters ***µ*** given the observed spike train *Y* is obtained by taking the logarithm of equation 16 and dividing by the number of spikes *N*:

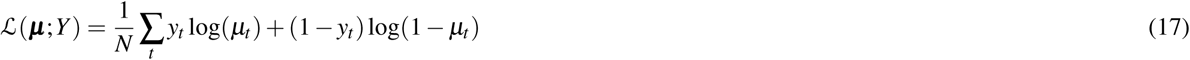

Adding the elastic net penalty,

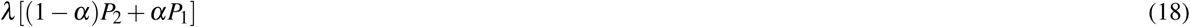

with

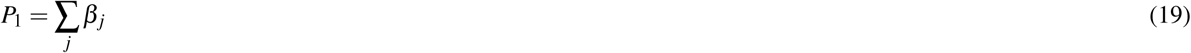

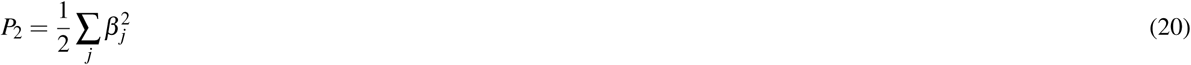

to the average negative log-likelihood function (Eq. 17) yields the loss function

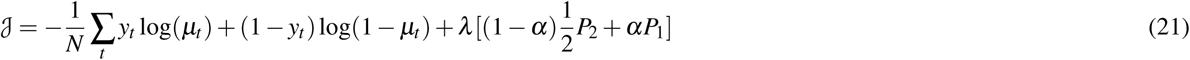

The values of the parameters *β*_0_ and ***β*** which minimize the loss function were found by batch gradient descent. The resulting ***β*** represents the linear filter of the LNP model estimated by maximum likelihood estimation (MLE). The hyperparameters *λ* (determining the strength of the regularization) and *α ∈* [0, 1] (determining the relative contributions of the L1 (Eq. 19) and L2 (Eq. 20) regularization terms) were determined by a cross-validated parameter grid search, choosing *λ* from 5 logarithmically spaced values from the interval [0.0001, 0.01], and *α* from {0, 0.01, 0.1, 0.5, 1}.

### Model prediction and evaluation

Both the STA fit and the MLE fit of the LNP were used to predict RGC spiking activity in response to the stimulus: the filter response of the stimulus was computed, and the nonlinearity was applied to the filter response, yielding the firing rate prediction. To prevent overfitting and overestimation of the model prediction performance, the model was fit on four seconds of stimulus, and the parameters derived from that fit were used to predict the firing rate in the remaining leftout second of the stimulus. This was done for all five possible splits into four training seconds and one test second.

Prediction performance *P* was evaluated as the correlation coefficient between true (*r*) and predicted firing rate 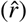 at a resolution of 1 kHz:

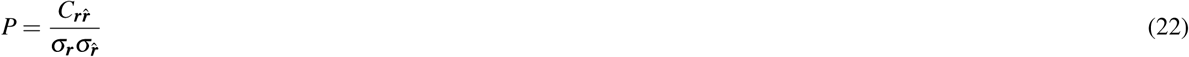

where *C* denotes the covariance matrix of the true and the estimated firing rates *r* and *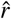*

### Hierarchical clustering

A hierarchical clustering algorithm was used to cluster the filters of all cells to determine whether there were different classes of filters. At first, the dimensionality of the filters was reduced from *N* = 300 features (30 ms at a sampling rate of 10 kHz) to *k < N* features retaining 90 % of the variance by applying principal component analysis (PCA) and projecting the filters onto the retained *k* principal components (PCs). Hierarchical clustering was then performed on these projections, using the `scipy.cluster.hierarchy` package with the average euclidean distance in PC space as clustering metric. The number of clusters was determined by visual inspection of the resulting dendrogram.

## Supplementary Information

**Figure S1.**
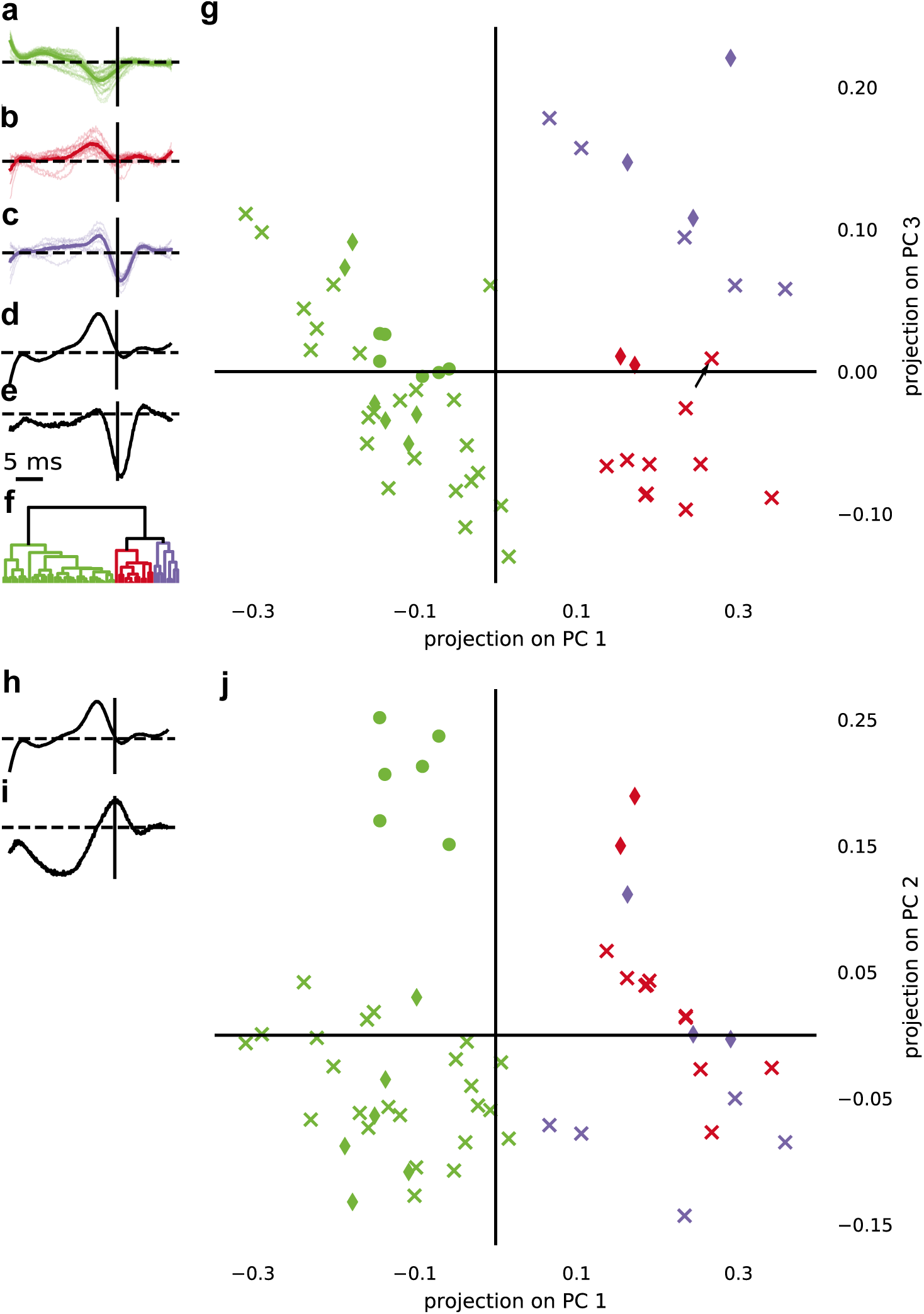
Hierarchical clustering of RGC electrical temporal filters. (a) - (c) Electrical temporal filters of all RGCs from wild-type retina recovered from the MLE fit of the LNP model, displayed separately for the three clusters identified by the hierarchical clustering algorithm. Thin lines are individual cell filters, thick lines indicate the average filter for one cluster. (d), 1st and 3rd principal component (PC) recovered from principal component analysis of the ensemble of temporal filters from all cells. (f) Dendrogram showing the separation of consecutively joined clusters along the clustering metric (distance in euclidean space). (g) Scatter plot of the projections of the temporal filters onto the 1st and 3rd PCs (shown in (d) and (e)). Colors indicate cluster assignment. Different markers (filled circle, cross and diamond) indicate cells recorded in different sessions. The black arrow marks the cell for which the assignment to clusters did not agree between STA and MLE estimate of the filters. (h) and (i) 1st and 2nd PC recovered from principal component analysis of the ensemble of temporal filter from all cells. (j) Scatter plot of the projections of the temporal filters onto the 1st and 2nd PCs (shown in (h) and (i)). Colors indicate cluster assignment. Different markers (filled circle, cross and diamond) indicate cells recorded in different sessions. Note that the 2nd PC separates filter projections of cells from different recordings, but does not separate the red and the violet cluster.

## Acknowledgements

This work was supported by the Baden-Württemberg Stiftung (NEU013), the German Ministry of Science and Education (BMBF, FKZ 01GQ1601 and FKZ 031L0059A) and the German Research Foundation through a Heisenberg Professorship (BE5601/4-1) and the SFB 1233 “Robust Vision” (project number 276693517).

## Author contributions statement

L.H., P.B. and G.Z. conceived the study. L.H. performed experiments and analyses. P.B. and G.Z. supervised the project. L.H. drafted the manuscript with input from PB and GZ. All authors reviewed the final manuscript.

## Additional information

### Availability of materials and data

The datasets generated and analysed during the current study, as well as the code used for analysis, are available from the corresponding author on reasonable request.

