## Supplemental Figure 1 for "Probing and predicting ganglion cell responses to smooth electrical stimulation in healthy and blind mouse retina"

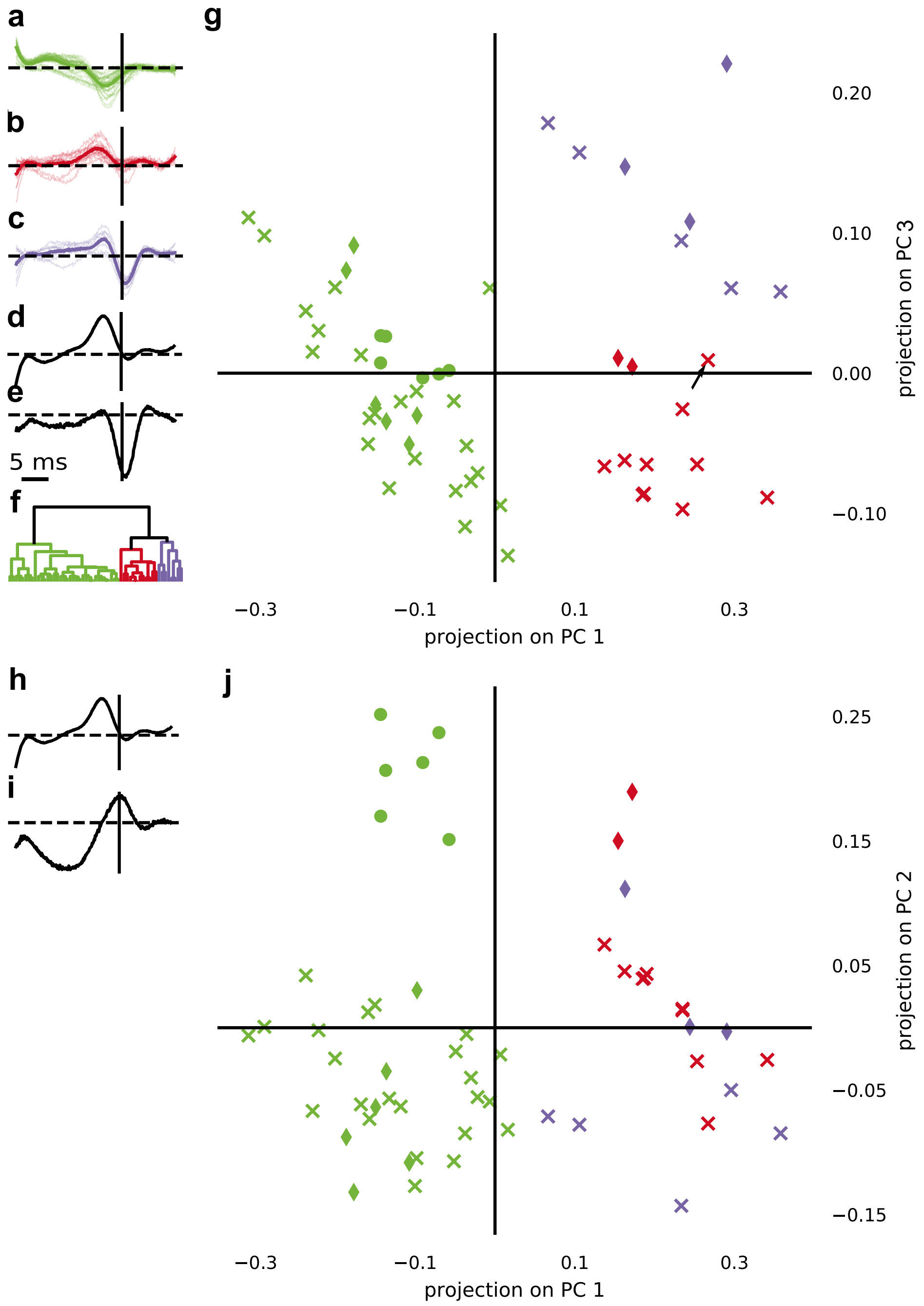


**Figure S1.** **Hierarchical clustering of RGC electrical temporal filters** (a) - (c) Electrical temporal filters of all RGCs from wild-type retina recovered from the MLE fit of the LNP model, displayed separately for the three clusters identified by the hierarchical clustering algorithm. Thin lines are individual cell filters, thick lines indicate the average filter for one cluster. (d), (e) 1st and 3rd principal component (PC) recovered from principal component analysis of the ensemble of temporal filters from all cells. (f) Dendrogram showing the separation of consecutively joined clusters along the clustering metric (distance in euclidean space). (g) Scatter plot of the projections of the temporal filters onto the 1st and 3rd PCs (shown in (d) and (e)). Colors indicate cluster assignment. Different markers (filled circle, cross and diamond) indicate cells recorded in different sessions. The black arrow marks the cell for which the assignment to clusters did not agree between STA and MLE estimate of the filters. (h) and (i) 1st and 2nd PC recovered from principal component analysis of the ensemble of temporal filter from all cells. (j) Scatter plot of the projections of the temporal filters onto the 1st and 2nd PCs (shown in (h) and (i)). Colors indicate cluster assignment. Different markers (filled circle, cross and diamond) indicate cells recorded in different sessions. Note that the 2nd PC separates filter projections of cells from different recordings, but does not separate the red and the violet cluster.
